# Perturbation of Lipid Bilayers by Biomimetic Photoswitches Based on Cyclocurcumin

**DOI:** 10.1101/2022.09.11.507454

**Authors:** Anastasiia Delova, Raúl Losantos, Jérémy Pecourneau, Yann Bernhard, Maxime Mourer, Andreea Pasc, Antonio Monari

## Abstract

The use of photoswitches which may be activated by suitable electromagnetic radiation is an attractive alternative to conventional photodynamic therapy. Here we report all atom molecular dynamics simulation of a biomimetic photoswitch derived from cyclocurcumin and experiencing *E/Z* photoisomerization. In particular, we show that the two isomers interact persistently with a lipid bilayer modeling a cellular membrane. Furthermore, the interaction with the membrane is strongly dependent on the concentration and a transition between ordered or disordered arrangements of the photoswitches is observed. We also confirm that the structural parameters of the bilayer are differently affected by the two isomers, and hence can be modulated through photoswitching, offering interesting perspectives for future applications.

**TOC Graphic:** 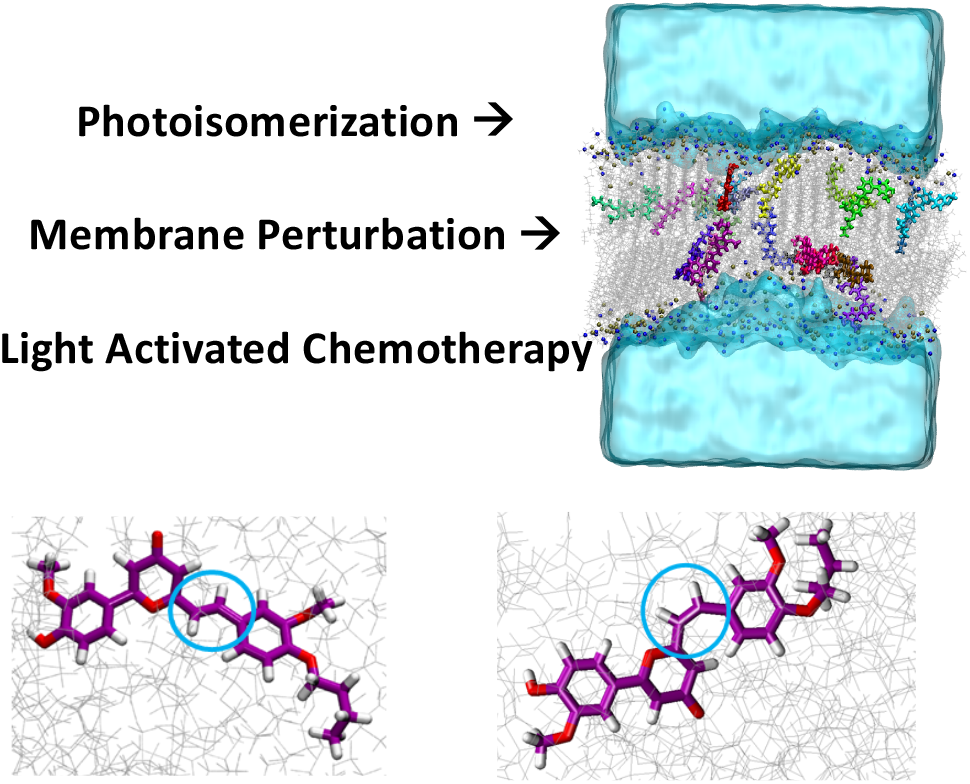

## INTRODUCTION

Conventional chemotherapy^1,2^ is based on the use of cytotoxic agents able to counteract the uncontrolled development of cancer cells, hence reducing tumor size and aggressivity, and should be counted among the major successes of modern medicine. As an example, *cis*-platinum,^3–5^ firstly proposed in the sixties is still widely used in conventional chemotherapy, especially against leukemia or testicular cancer.^6^ However, if it allows to treat a large number of malignancies and significantly increases the life expectancy, chemotherapy also suffers from crucial lack of selectivity, which correlates with heavy secondary effects, specially aggravated by the use of metals which in some cases are poorly metabolized and accumulated in the body.^7^ An attractive alternative to conventional chemotherapy, relies on the use of drug which are activated by a luminous stimulus. In this context, the most common strategy is the so-called photodynamic therapy (PDT)^8–11^ in which after light activation a sensitizer (the drug) activates molecular oxygen to its singlet state (^1^O_2_). The high reactive ^1^O_2_ will react with biological components, including cellular membranes, leading to the cell death by apoptosis or necrosis. While selectivity is achieved through the controlled use of the light source, form a photophysical point of view oxygen activation requires the population of the sensitizer’s triplet state *via* an intersystem crossing (ISC). PDT has nowadays found relatively widespread clinical applications in the treatment of different cancer types, as well as for non-malignant conditions such as psoriasis.^12^ Up to date, the most commonly used PDT agents are based on porphyrin or phthalocyanines sensitizers,^13,14^ which combine efficient and tunable ISC^15^ with absorption in the red or infrared wavelengths region, being necessary to guarantee a sufficient penetration of the light into the biological tissues. Recently, the use of nanoparticles,^16,17^ likewise exploiting plasmonic effect,^18^ has been proposed. Even if the photophysical and optical properties are optimal, PDT photosensitizers still suffers from a poor bioavailability, scarce water solubility and aggregation induced excited state quenching, which can be addressed by specific drug delivery strategies.^19–22^ Furthermore, the action mechanism of PDT requires a high availability of oxygen, which makes its use problematic in hypoxic conditions, commonly developed by some solid tumors.^23–25^ Therefore, alternative strategies, still based on the light activation of specific drugs have been designed, which can be grouped under the general name of light activated chemotherapy (LAC).^26^ In LAC, photochemical and photophysical pathways different from the oxygen activation are exploited to disrupt the biological macrosystems and induce the cellular death. As an example, the presence of suitable photoactivated leaving group in DNA interacting sensitizers^27,28^ has been proposed as a way to induce covalent interstrand linkages, while Ruthenium complexes have been proposed for the controlled photorelease of reactive species.^29,30^

In this respect a particular sounding possibility is offered by molecular photoswitches,^31–33^ which after photoisomerization may experience important structural reorganization, such as double bond isomerization. Indeed, if a persistent interaction between the photoswitch and a biological structure, such as a lipid cellular membrane exists, the light induced perturbation may be transmitted to the biological structure significantly altering its structural properties, and potentially leading to the triggering of death induction signals.^34,35^ Recently photoswitches have also been proposed as potential light-controlled regulators of serotonin transporters^36,37^ and ion channels as well for potential optogenetics^38–40^ applications. In previous contributions we have presented the synthesis and characterization of different biomimetic cyclocurcumin (CC)-based photoswitches,^41,42^ which can potentially be used in LAC framework. Indeed, CC is a natural and non-toxic component of turmeric, and through it possess a photoisomerizable double bond the isomerization quantum yield is strongly dependent on the solvent polarity.^43,44^ In addition, turmeric-based extract are also currently considered for their anticancer capacities, ^45–49^ including PDT.^50–53^ We have also shown that by introducing a g-pyrone analogous not only the absorption wavelength is red-shifted, as well the isomerization solvent-dependency is strongly reduced.^41,42^ Finally, by using a combination of experimental and modeling techniques, including enhanced sampling and free energy calculations, we have shown that appending the g-pyrone derivative with a hydrophobic butyl chain (CCBu, Figure 1) leads to the spontaneous penetration inside the lipid bilayer, and more specifically at the interface between the polar heads and the fatty acid core.^54^ Interestingly, the free energy profile for the insertion into the bilayer shows important differences between the *E* and the *Z* isomers, with the former developing a more favorable interaction with the lipid bilayer. Molecular modeling and simulation have also been complemented by biophysical determinations on liposomes and lipid monolayers confirming the perturbation induced after light absorption, and the differential effects of the two isomers.

**Figure 1.**
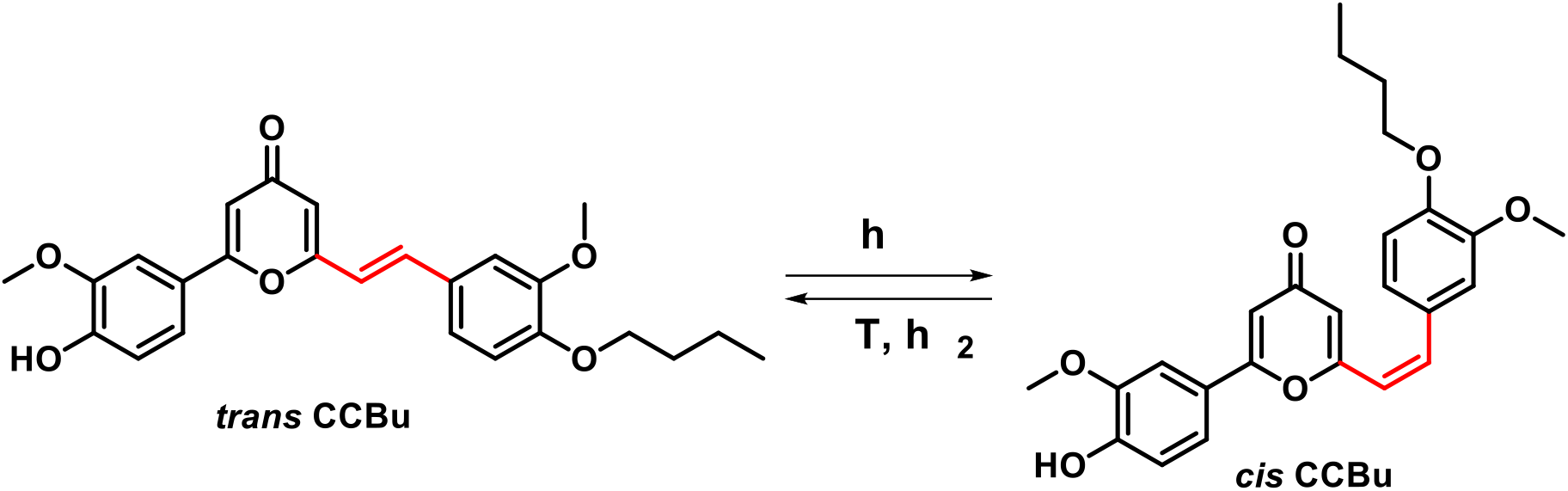
Schematic representation of the two isomers of the biomimetic analogous CCBu. The isomerizable double bond is highlighted in red.

In this contribution we want to use all atom molecular dynamic (MD) simulations to further unravel the atomistic details of the interactions taking place between CCBu and a model lipid bilayer. Particularly we aim to elucidate the effects due to the chromophore concentration and the interplay with the lipid membrane organization, also considering the possible self-aggregation of the chromophore. A particular attention will be devoted to unraveling the perturbation induced on the lipid membrane organization by the isomerization of CCBu as unraveled by the change in its structural parameters, through out of equilibrium MD simulations. Our results evidence that the interaction between CCBu and the lipid bilayer is, indeed, more complex than expected and ultimately give rise to a transition between ordered and disordered arrangements which are reflected on the different structural properties of the membrane. Our results also allow to pinpoint possible structural modifications, which could enhance the chromophore-induced perturbations of the bilayer, and at the same time they also permit to identify the optimal range of concentration to maximize the effects due to the switching.

## COMPUTATIONAL METHODOLOGY

### General MD Simulations

All the MD simulations have been performed using the NAMD code^55,56^ and have been analyzed and visualized using VMD^57^ and its Membrane Plugin extension.^58^ For all the simulations CCBu has been described using the General Amber Force Field (GAFF) approach.^59^ Atomic point charges have been obtained through the fitting of the restricted electrostatic potential (RESP) procedure. Coherently with the GAFF protocol, and with our previous works,^41–43^ the electrostatic procedure has been obtained from the Hartree-Fock wavefunction obtained with the 6-31G* basis set. Note that the ground state geometry of the chromophore in its *E* and *Z* configuration was previously optimized at Density Functional Theory (DFT) level using B3LYP functional and the 6-31+G* basis set.^41,42^ All the quantum chemistry calculations have been performed using the Gaussian 16 set of codes.^60^ Noteworthily, two independent force fields have been parameterized for the *E* and the *Z* isomer, respectively. In addition to different concentrations of *E* or *Z* CCBu the simulation box contains a lipid bilayer composed of 159 dipalmitoylphosphatidylcholine (DPPC) lipids on each leaflet. A water buffer of 35 Å has been added including K^+^ and Cl^−^ ions to mimic the physiological salt concentration of 0.15 M. DPPC have been described through the Amber14^61^ lipid force field, while water is modeled with TIP3P.^62^ All the initial systems have been prepared using the CHARMM-GUI web interface^63^ and hydrogen mass repartition (HMR)^64^ is systematically applied allowing the use of a 4.0 fs time step to numerically solve the Newton equations of motion. Each initial system has been subjected to 50000 minimization steps *via* the conjugated gradient algorithms, followed by equilibration and thermalization by progressively remove positional restraints on the lipid heavy atoms during 9 ×10^6^ steps. All the MD simulation have performed in the isothermal and isobaric ensemble (NPT) at 300 K and 1 atm by using Langevin thermostat^65^ and barostat.^66^ Initial systems have been constructed including one, two, five, ten, fifteen, and twenty molecules of *E*-CCBu placing it/them in the water bulk and MD simulation of 660 ns have been performed for each system, the stability is being checked through the analysis of the root mean square deviation (RMSD) evolution. The chosen initial conditions correspond to a CCBu/lipid ratio of 0.63%, 1.26%, 3.14%, 6.29%, 9.43%, and 12.58%, respectively. By taking into account the size of the box the previous values can also be translated in CCBu molar concentration yielding 1.7 10^−6^, 3.3 10^−6^, 8.3 10^−6^, 1.8 10^−5^, 2.5 10^−6^, 3.3 10^−5^ M, respectively.

Note that equilibrium MD simulation was also performed on the lipid bilayer without any CCBu as a control.

To better understand the effect of different concentrations of CCBu on the membrane structure during the dynamics we have calculated the averaged values of the area per lipid^58^ along the trajectory. Furthermore, we have estimated the membrane thickness by calculating the lipid interdigitation index.^58^ Finally, the deuterium order parameter (-S_CD_)^67^ providing information on the flexibility and disorder of the different carbon atoms composing the fatty acid chains has been obtained.

### Free energy profiles

To take into account the effects due to aggregation of the chromophores we have calculated the free energy profiles for the dissociation of a *E* CCBu dimer in a water box, monitoring the distance between the center of mass of each chromophore as collective variable. Furthermore, we also obtained the free energy cost due to the penetration of a *E* CCBu dimer in the DPPC bilayer, which was explored by monitoring the distance between the center of mass of the CCBu dimer and the center of the membrane projected on the membrane axis (z). In this case to ensure that the dimer does not dissociate during the penetration through the membrane a harmonic constraint on the distance between the center of masses of each CCBu monomer is applied keeping it around the equilibrium intermonomer distance. All the free energy calculations have been performed using the combination of metadynamics^68^ and extended adaptative biased force^69^ (meta-eABF)^70,71^ as implemented in Colvar^72^ and NAMD.

### Steered Molecular Dynamics (SMD)

To allow the inclusion of multiple molecules of CCBu in our model lipid bilayer we performed steered molecular dynamics (SMD)^73^ starting from an initial conformation in which the chromophores are placed in the water bulk. Systems consisting of one, five, ten, fifteen, and twenty CCBu monomers in their *E* conformation were built. SMD was realized applying a harmonic force of 10 kcal/mol over 100,000 steps on top of a collective variable consisting of the z, *i.e.* the membrane axis, component of the distance between the center of mass of the monomers and the center of the membrane. After SMD, equilibrium MD simulation was run for 660 ns, allowing to obtain the average membrane structural parameters as previously detailed (Figure 2A).

**Figure 2.**
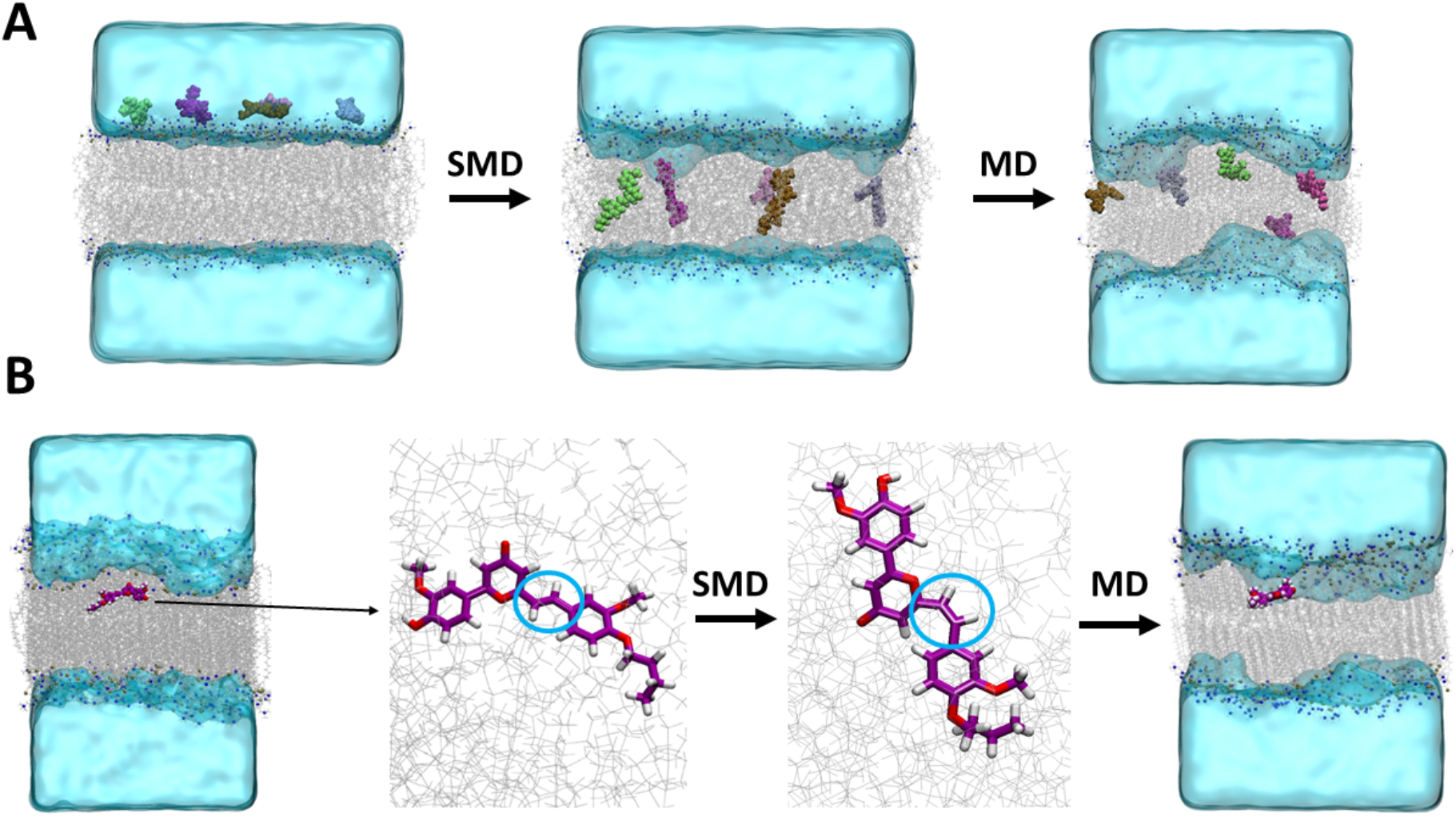
Schematic representation of the SMD protocol for the penetration of multiple CCBu molecules into the lipid bilayer (A) and for the simulation of the E to Z photoisomerization effects (B).

SMD was also used to simulate the fast, compared to the inherent membrane degrees of freedom, isomerization of CCBu. To this end, starting from the equilibrated systems comprising CCBu embedded in the lipid bilayer, we applied a harmonic constraint of 5 kcal/mol to the dihedral angle around the isomerizable double bond for 100 000 steps. After obtaining a collection of *Z* isomers embedded in the bilayer, the systems were allowed to relax through MD simulations whose length reaches 660 ns for each system, and the membrane structural parameters have been calculated again (Figure 2B). In the case of the system composed of 5 chromophores embedded in the lipid bilayer we also considered the effects of an incomplete isomerization by constructing initial situation in which the ratio between E/Z isomers varied from 0/5 to 5/0.

## RESULTS AND DISCUSSSION

In a previous contribution we observed that a single CCBu molecule in *E* conformation is indeed spontaneously entering a DPPC lipid bilayer to accommodate at the interface between the lipid polar heads and the hydrophobic core.^54^ These results have also been confirmed by the determination of the free energy profile for the penetration of one *E* CCBu molecule which has been found to be barrierless. However, when adding more than one monomer a more complicated scenario is operating. Unsurprisingly, due to the relatively highly hydrophobic nature of CCBu, self-aggregation takes place at moderate concentration of the chromophore, as can be clearly observed for the systems containing five and ten *E* CCBu (Figure 3). Obviously, the large size of the aggregate precludes its spontaneous penetration on the lipid bilayer. However, a more complex behavior is observed for the system containing two *E* CCBu moieties, in this case, indeed, we observed the penetration of one monomer, while the second one remains in the water bulk (Figure 3). All these results suggest a rather complex interplay and equilibrium between aggregation and membrane penetration.

**Figure 3.**
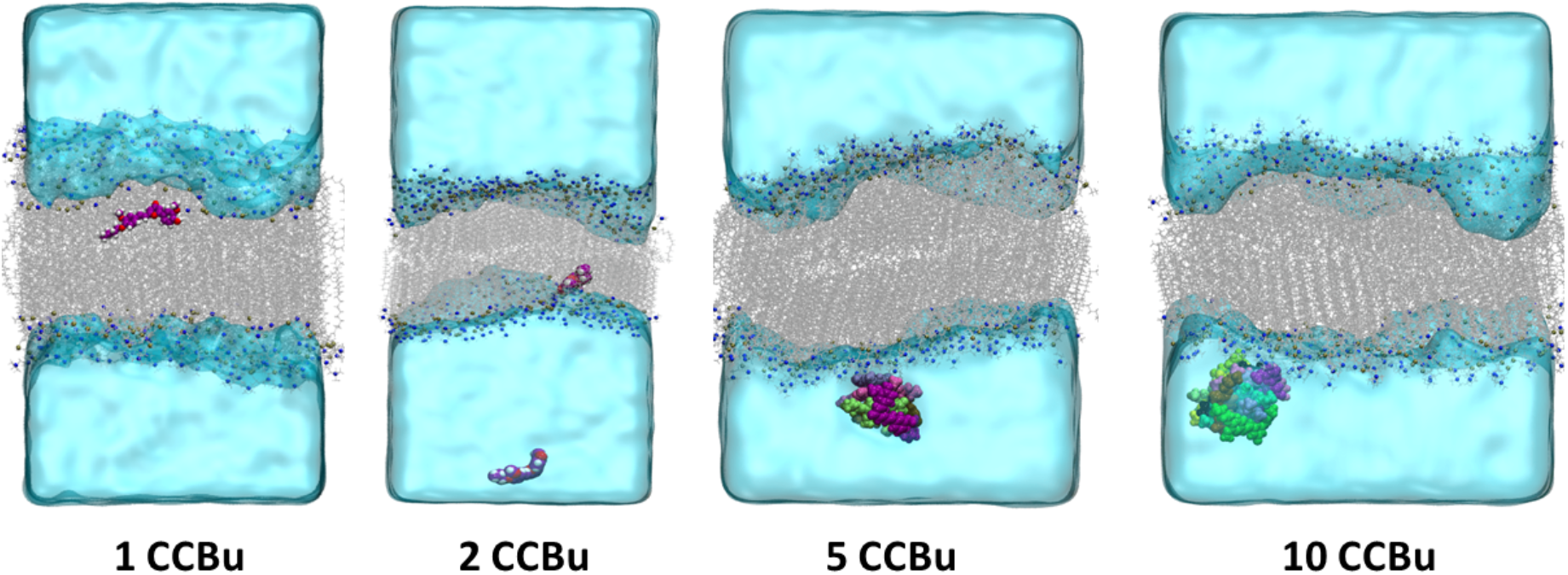
Representative snapshots for the MD simulations on the systems containing different CCBu and showing the self-aggregation at high concentration.

To disentangle theses effects, we calculated the free energy profile for the separation of a CCBu dimer in water solution and for the penetration of the same dimer inside the lipid bilayer, which are presented in Figure 4. We may see that the dimer is indeed stable, however its dissociation free energy is of *c.a.* 4.5 kcal/mol, which is coherent with the dominant dispersion and π-stacking interactions which are developed. Although, the dissociation free energy of the dimer and an aggregate could differ they will likely remain of the same order of magnitude. To confirm this hypothesis, we report in ESI (Figure S5) the free energy profile for the dissociation of one CCBu unit from a 5-member aggregate, which yields a penalty of about 4.0 kcal/mol, i.e. significantly close to the one of the dimer.

**Figure 4.**
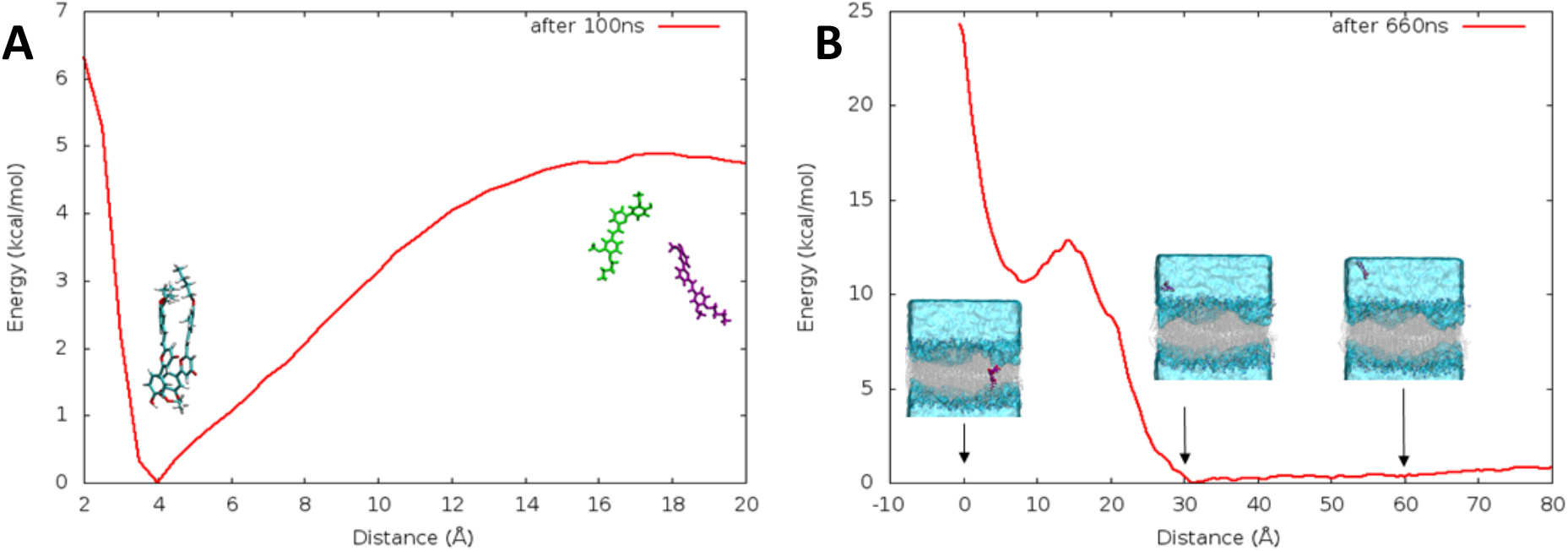
Free energy profile for the dissociation of the CCBu dimer in water solution (A) and for the penetration of a constrained CCBu dimer in the DPPC bilayer. Representative snapshots extracted through the collective variables are also provided as inlays.

On the other hand, the penetration of an undissociated dimer in the DPPC membrane is accompanied by a relatively high free energy penalty and would necessitate overcoming more than 12 kcal/mol in correspondence to the insertion in the polar head region. Furthermore, only a metastable state can be reached for the constrained dimer embedded on the membrane. These results are not unexpected and coupled with the previously determined free energy profile for the penetration of a single monomer, which is barrierless and associated to a stabilization of about 5.0 kcal/mol compared to the water bulk, allowing to rationalize the observation of the equilibrium MD simulations. Indeed, while the penetration of larger aggregates is clearly thermodynamically and kinetically unfavorable, the relatively small dissociation energy of the dimer and the stabilization of the monomer in the bilayer suggests that while the aggregates are forming in water solution they could dissociate at the contact with the membrane allowing the penetration of a single monomer at once. However, we have not observed this occurrence through our MD simulations. This is not surprising since considering the energy penalties into play the event is still rare at the time-scales of our equilibrium MD, i.e. hundreds of nanoseconds. Instead, it should take place in the millisecond time-range, i.e. still relevant for chemical or biological applications.

Thus, we further studied the behavior of our model lipid model in presence of increasing concentration of *E* CCBu embedded in the membrane. As detailed in the Methodological Section to enforce the loading of the chromophore inside the membrane we have resorted to an SMD approach, after which the bilayer and the chromophores are allowed to mutually equilibrate without restrains. Representative snapshots of the membrane loaded with different concentrations of CCBu are reported in Figure 5 (other snapshots are presented in the SI). Notably, after equilibration we never observed any evidence of spontaneous release of CCBu back to the water bulk, even for the larger concentrations studied, thus indicating a rather persistent interaction between the chromophore and the model membrane. Already from the visual inspection of the MD trajectories we can evidence a distinctive concentration-dependent organization of CCBu. Indeed, at low concentration, *i.e.* up to five chromophores, the ligands are occupying the polar head region with a relatively high degree of disorder. Indeed, the coexistence of conformations in which the main axis of *E* CCBu is either parallel or orthogonal to the polar heads. However, when increasing the chromophore concentration, we reached an intermediate region in which *E* CCBu acquires a higher degree of order to maximize space occupancy. Indeed, in this case, we may observe a prevalence of configurations parallel to the membrane axis, which may also be stabilized by π-stacking. This ordered arrangement cannot be sustained when further increasing the concentration of chromophore, and indeed for a loading of twenty, we observe again a rather disordered arrangement in which *E* CCBu occupy all the available space in the confinement environment of the membrane. Thus, we may see that a transition between three arrangements takes place depending on the chromophore loading and concentration. Indeed, after a first disordered disposition is established at low concentration, in the intermediate concentration range an ordered arrangement is observed, which is however broken at high concentrations to reach a novel disordered situation.

**Figure 5.**
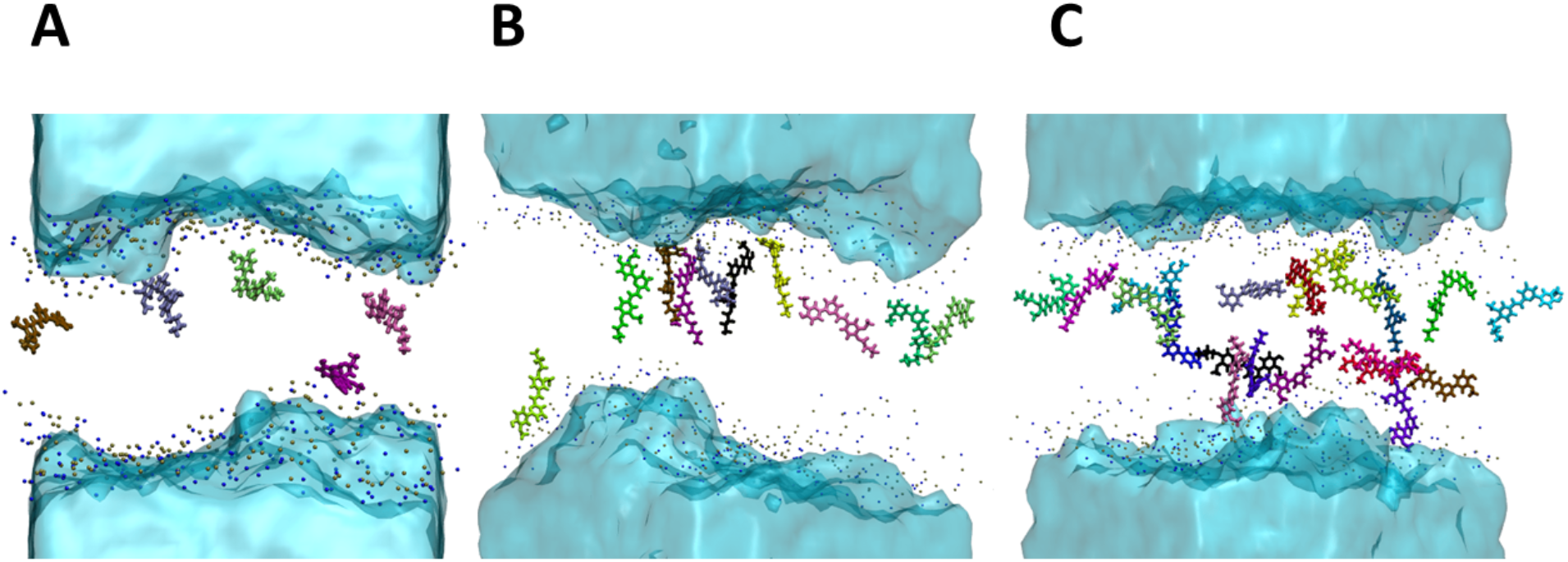
Representative snapshots of the MD simulations for the systems containing five (A), ten (B), and 20 (C) *Z* CCBu chromophores embedded inside the lipid bilayer.

As shown in Figure 6, after simulating the *E* to *Z* photoisomerization of CCBu at various concentrations by forcing the rotation around the isomerizable double bond by SMD, we still observe a persistent interaction with the lipid bilayers. Indeed, once again no spontaneous release of any *Z* isomer is observed during the time laps covered by our simulations. Representative configurations illustrating the behavior of the lipid membrane in presence of *Z* CCBu are reported in Figure 6 and in ESI. Furthermore, and interestingly, we may observe that three arrangements observed previously are conserved even after isomerization. Certainly, while at low concentration a disordered arrangement is observed, at intermediate range we may observe that, despite the lack of a rigid and planar core, *Z* CCBus assume a more ordered arrangement which is still roughly parallel to the membrane axis and driven by π-stacking interactions. Finally, at higher concentrations a disordered arrangement is achieved, again most probably to maximize the occupancy of the confined environment, however the more compact and less extended structure of the Z isomer seems favorable to maintain more ordered arrangements.

**Figure 6.**
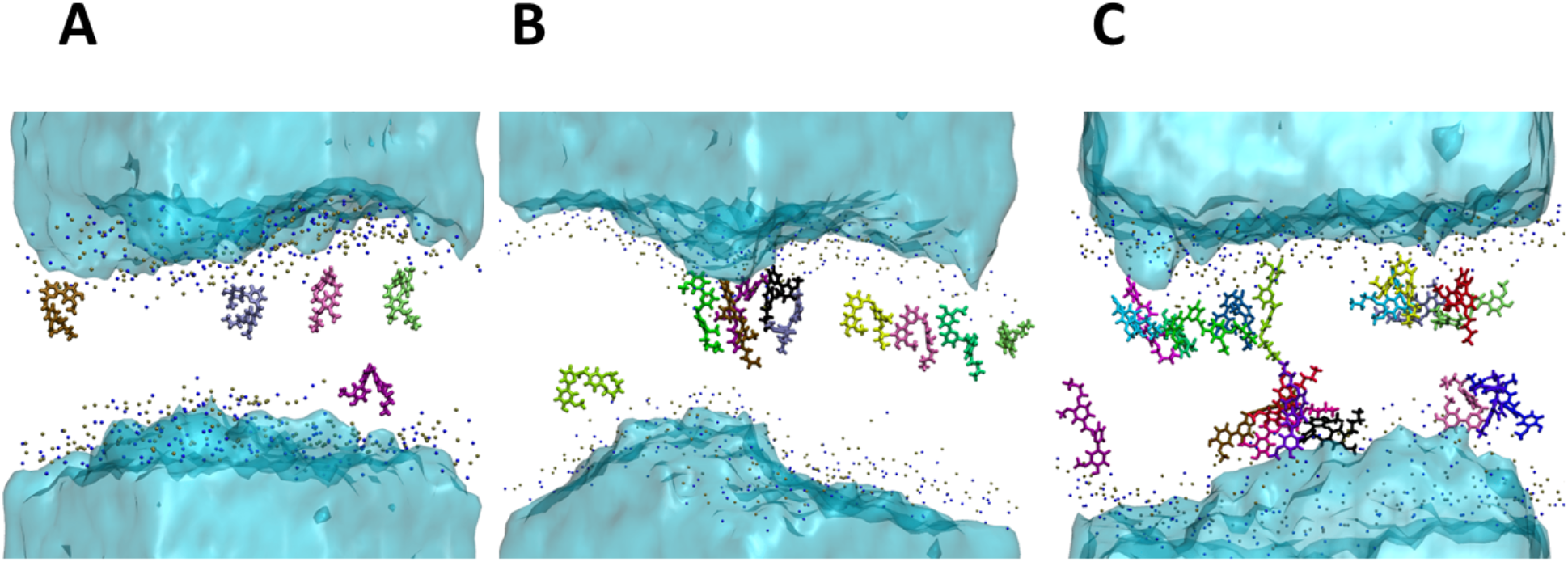
Representative snapshots of the MD simulations for the systems containing five (A), ten (B), and 20 (C) *Z* CCBu chromophores embedded inside the lipid bilayer.

The effects of the perturbation observed in the arrangement of the CCBu chromophores is also reflected and transferred to the global structural properties of the lipid membrane and in particular the area per lipid and the lipid interdigitation (Figure 7) which clearly present a non-linear behavior. Indeed, the value of the area per lipid increases from 0 to 5 *E* CCBus embedded, reaching values which, albeit relatively small, are significantly different from those of the control membrane. Interestingly for each point of the low concentration area the perturbation induced by *Z* CCBu is larger than for the *E* isomer. The difference between the effects of the two isomers increased with the concentration and reaches its maximum for a loading of 5 chromophores. Pointing that by photoinduced isomerization a larger destabilization of the biological membrane can be achieved. The values of the lipid interdigitation also increase steadily for this concentration domain, indicating that the thickness of the membrane is decreasing, while the *Z* isomers is still causing a much stronger structural perturbation. However, in the intermediate concentration range (ten and fifteen chromophores) we observe a sharp decrease of both, the area per lipid and the lipid interdigitation parameters which drops back to values very close to those of the unperturbed membrane. Interestingly, increasing the number of embedded chromophores in this concentration domain leads to a further decrease of both structural parameters, *i.e.* a behavior opposite to the one observed at low concentrations. In the intermediate regime the difference in the structural parameters for *Z* and *E* CCBu embedded bilayer is negligible and below the error bars, hence in this regime, and most probably due to the ordered arrangement which is maintained in both cases no net effect of the isomerization can be pinpointed. Finally, when reaching the high concentration phase (*i.e.* twenty embedded chromophores) we observe once more a sharp increase of the average values of both structural parameters and particularly the area per lipid. This fact is due to the formation of a novel disordered phase, being more pronounced for the *E* than the *Z* isomer, and different from the low concentration regime. This difference may arise from the more extended and more rigid *E* isomers which is assuming a less compact arrangement compared to the more globular *Z* isomers. Indeed, as also shown in ESI the loading of the lipid bilayer with twenty *E* CCBus has a very sharp effect on the Deuterium order parameter, while the effect is much smoother in the case of the *Z* isomer.

**Figure 7.**
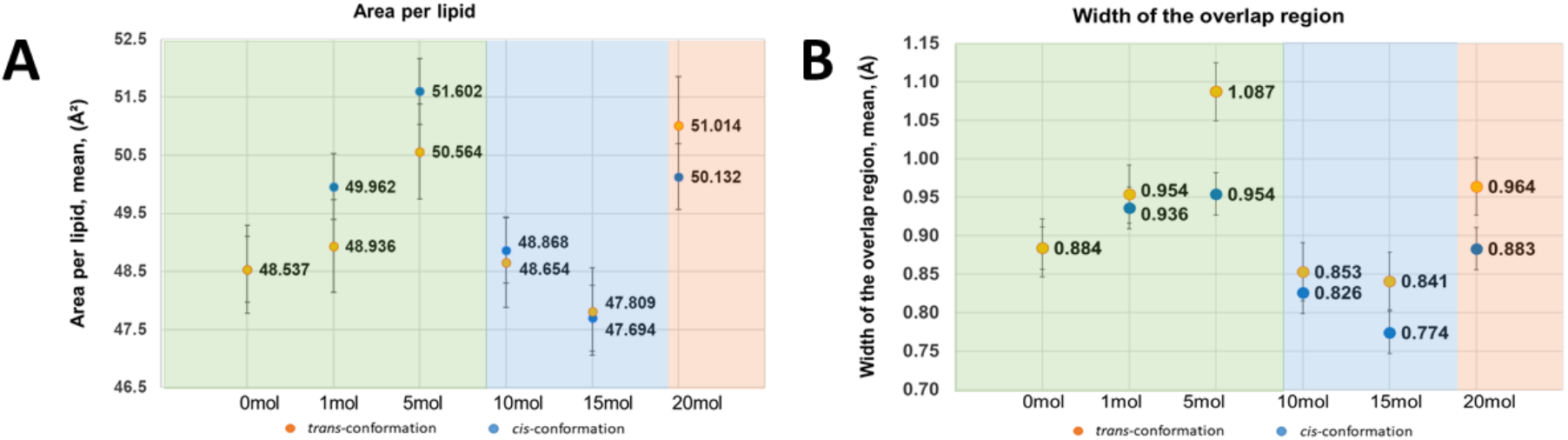
Average values calculated along the MD simulations for the area per lipid (A) and the lipid interdigitation (B) as a function of different concentrations of *E* and *Z* CCBu, note that the standard deviations are also reported. The three ordered/disordered/ordered arrangements are also highlighted by the different colors of the background in panel A and B.

It has to be pointed out that in the present study we have fully considered only the isomerization of whole the chromophores embedded in the lipid bilayer. This represent clearly an approximation, since the isomerization quantum yield would not be unitary. However, the study of partially isomerized systems will be highly complex, and would require a combinatorially high number of starting structures. However, to at least estimate the effects of a partial isomerization we report in ESI (Figure S4) the average values of the area per lipid obtained while changing the E/Z ratio for 5 chromophores embedded in the bilayer from 0/5 to 5/0. These results, even if preliminary clearly indicate that significantly difference appears already for 2/3 and 3/2 ratios.

The presence of three different and well defined arrangements, and the differential perturbation induced on the lipid bilayer properties, not only confirms the possible utilization of our chromophores as light activated cellular membrane perturbators, but they also allow to identify an optimal concentration range to maximize the effects induced by the photoisomerization, which should correspond to the highest concentrations supporting the first disordered arrangement (in our simulations corresponding to 5 *E* or *Z* CCBu). Indeed, for these values the difference in the structural parameters in presence of the two isomers is maximal. The high concentration regime would probably be difficult to achieve in biological relevant conditions, and it would require loading the membrane with the thermodynamically unfavored Z isomer.

## CONCLUSIONS

In this contribution we have deeply characterized the behavior of biomimetic cyclocurcumin photoswitches, CCBu, considering their capacity to interact with a model lipid bilayer. In particular we have confirmed that while the chromophores tend to aggregate in water solution their stabilization energy is moderate, and the aggregate can be dissolved at the membrane interface to allow the loading of one drug at the time. More importantly, we have also observed that the interaction of CCBu leads to the formation of three distinctive concentration dependent arrangements: a disordered disposition at low concentration, followed by an ordered one at intermediate range and a novel disordered disposition at high concentration. We have also shown that the structural perturbation induced on the lipid membrane in the low concentration regime are higher for the *Z* than the *E* isomer. This situation is reversed at high concentration, where the *E* isomer is more destabilizing, while in the intermediate range no difference between the two isomers can be observed. Thus, the highest possible concentration of the low regime appears as the most favorable for inducing a biological relevant effect of the photoswitching. Although, the structural differences are significant, *i.e.* above the errors bar, the perturbations of the membrane still appear moderate and could be increased for instance by appending bulky and/or rigid groups to the switch. However, if this strategy is attractive care should be taken that the proposed substitution would not result in too high energy penalty for the internalization of the chromophores, or to a viscosity-impeded isomerization. To minimize these risks, probably a combination of hydrophobic and amphiphilic groups should be considered. Thus, in the following we plan to explore this path by studying modified CCBu with the same protocol used in the present contribution, also to foster a rationally design development of novel agents for LAC. Furthermore, we also plan to partially close the gap with the biological systems by complexifying our membrane model to include different lipid compositions, as well as cholesterol, whose influence on the structural properties, and in particular the bilayer rigidity, can be remarkable.

## DATA and SOFTWARE AVAILABILITY

NAMD and VMD are available free of charges for academic users. All the MD trajectories are stored in the University Computer center and are available upon request.

## ASSOCIATED CONTENT

### Supporting Information

Coordinates and point charges of the force field for *Z* and *E* CCBu, RMSD time series for all the MD simulations, snapshots of other chromophores concentrations in both Z and E conformations, −S_CD_ parameters for *Z* and *E* isomers. The following files are available free of charge. (PDF)

### Supporting Information

Coordinates

## AUTHOR INFORMATION

### Author Contributions

The manuscript was written through contributions of all authors. All authors have given approval to the final version of the manuscript.

## ACKNOWLEDGMENT

The authors thank GENCI and Explor computing centers for computational resources. A.D. thanks Université Paris Cité for her Ph.D. fellowships. R.L. thanks Universidad de La Rioja and Ministerio de Universidades for his “Margarita Salas” grant. Support from the European Union European Regional Development Funds (Programme opérationnel FEDER-FSE Lorraine et Massif des Vosges 2014-2020 “Fire Light” project: “Photo-bio-active molecules and nanoparticles”) is also acknowledged. A.M. thanks ANR and CGI for their financial support of this work through Labex SEAM ANR 11 LABX 086, ANR 11 IDEX 05 02. The support of the IdEx “Université Paris 2019” ANR-18-IDEX-0001 and of the Platform P3MB is gratefully acknowledged.

