## Supplementary Information for "Perturbation of Lipid Bilayers by Biomimetic Photoswitches Based on Cyclocurcumin"

**Potential of Mean Force for the dissociation of one CCBu unit from a 5-membered aggregate** .. 14

### Cartesian coordinates and charges of *trans* and *cis*-CCBu

To facilitate the visualization the mol2 files of both isomers were included, also providing the QM charges used for the needed parametrization following the GAFF strategy. These QM charges were obtained under response calculation at HF/6-31G* level of theory.

***trans*-CCBu**

```
@<TRIPOS>MOLECULE
CBU
  57  59  1  0  0
SMALL
resp

@<TRIPOS>ATOM
   1 C1     -5.0030  -2.7680  -0.7210 ca    1 CBU   -0.239554
   2 C2     -6.3540  -2.6860  -0.4050 ca    1 CBU    0.272242
   3 C3     -6.8880  -1.4600   0.0500 ca    1 CBU    0.190995
   4 C4     -6.0750  -0.3450   0.1800 ca    1 CBU   -0.225953
   5 C5     -4.7050  -0.4230  -0.1500 ca    1 CBU   -0.114296
   6 C6     -4.1840  -1.6460  -0.5930 ca    1 CBU   -0.174799
   7 C7     -3.8480   0.7680  -0.0360 cc    1 CBU    0.493676
   8 O1     -2.5180   0.4470   0.0000 os    1 CBU   -0.311238
   9 C8     -1.5620   1.4290   0.0990 cc    1 CBU    0.483006
  10 C9     -1.9120   2.7380   0.1910 cd    1 CBU   -0.670893
  11 C10    -3.3060   3.1680   0.1510 c     1 CBU    0.901030
  12 C11    -4.2540   2.0610   0.0180 cd    1 CBU   -0.681997
  13 O2     -3.6550   4.3500   0.2190 o     1 CBU   -0.645101
  14 C12     2.2820   0.9960   0.0220 ca    1 CBU    0.048624
  15 O3     -7.1570  -3.7750  -0.5240 oh    1 CBU   -0.563218
  16 O4     -8.2280  -1.5220   0.3410 os    1 CBU   -0.293945
  17 C13     3.3580   1.8900  -0.1870 ca    1 CBU   -0.314286
  18 C14     4.6780   1.4560  -0.1930 ca    1 CBU    0.258286
  19 C15     4.9610   0.0810   0.0180 ca    1 CBU    0.330382
  20 C16     3.9020  -0.8050   0.2300 ca    1 CBU   -0.283542
  21 C17     2.5820  -0.3550   0.2330 ca    1 CBU   -0.221286
  22 O5      5.7650   2.2530  -0.3890 os    1 CBU   -0.336663
  23 O6      6.2750  -0.2590  -0.0020 os    1 CBU   -0.474312
  24 C18    -8.8750  -0.3420   0.7980 c3    1 CBU   -0.125739
  25 C19     5.5460   3.6370  -0.6040 c3    1 CBU    0.026521
  26 C20    -0.2310   0.8410   0.1030 ce    1 CBU   -0.342579
  27 C21     0.9250   1.5330   0.0060 cf    1 CBU   -0.038913
  28 H1     -4.6060  -3.7160  -1.0690 ha    1 CBU    0.194953
```

```
   29 H2         -6.4840    0.5840    0.5570 ha        1 CBU       0.166619
   30 H3         -3.1330   -1.7180   -0.8480 ha        1 CBU       0.164719
   31 H4         -1.1500    3.5020    0.3000 ha        1 CBU       0.226795
   32 H5         -5.3050    2.3120   -0.0510 ha        1 CBU       0.227206
   33 H6         -8.0510   -3.5000   -0.2550 ho        1 CBU       0.409175
   34 H7          3.1360    2.9390   -0.3490 ha        1 CBU       0.173244
   35 H8          4.1030   -1.8560    0.3980 ha        1 CBU       0.149378
   36 H9          1.7870   -1.0730    0.4100 ha        1 CBU       0.197761
   37 H10        -9.9210   -0.6110    0.9510 h1        1 CBU       0.107490
   38 H11        -8.4440    0.0040    1.7460 h1        1 CBU       0.107490
   39 H12        -8.8080    0.4600    0.0530 h1        1 CBU       0.107490
   40 H13         6.5350    4.0780   -0.7350 h1        1 CBU       0.063969
   41 H14         4.9460    3.8160   -1.5070 h1        1 CBU       0.063969
   42 H15         5.0500    4.1070    0.2560 h1        1 CBU       0.063969
   43 H16        -0.2240   -0.2430    0.1780 ha        1 CBU       0.158621
   44 H17         0.8660    2.6150   -0.1080 ha        1 CBU       0.141473
   45 C22         6.6350   -1.6250    0.2010 c3        1 CBU       0.306458
   46 H18         6.2710   -1.9650    1.1820 h1        1 CBU       0.001941
   47 H19         6.1640   -2.2530   -0.5690 h1        1 CBU       0.001941
   48 C23         8.1520   -1.7200    0.1240 c3        1 CBU      -0.112489
   49 H20         8.4780   -1.3390   -0.8520 hc        1 CBU       0.055166
   50 H21         8.5840   -1.0530    0.8810 hc        1 CBU       0.055166
   51 C24         8.6630   -3.1520    0.3290 c3        1 CBU       0.096147
   52 H22         8.2150   -3.8130   -0.4270 hc        1 CBU      -0.015522
   53 H23         8.3210   -3.5270    1.3040 hc        1 CBU      -0.015522
   54 C25        10.1900   -3.2530    0.2520 c3        1 CBU      -0.175700
   55 H24        10.5280   -4.2840    0.4010 hc        1 CBU       0.043882
   56 H25        10.5590   -2.9170   -0.7250 hc        1 CBU       0.043882
   57 H26        10.6660   -2.6300    1.0180 hc        1 CBU       0.043882

@<TRIPOS>BOND
    1    1    2 ar
    2    1    6 ar
    3    1   28 1
    4    2    3 ar
    5    2   15 1
    6    3    4 ar
    7    3   16 1
    8    4    5 ar
    9    4   29 1
   10    5    6 ar
   11    5    7 1
```

| | | | |
|---|---|---|---|
| 12 | 6 | 30 | 1 |
| 13 | 7 | 8 | 1 |
| 14 | 7 | 12 | 2 |
| 15 | 8 | 9 | 1 |
| 16 | 9 | 10 | 2 |
| 17 | 9 | 26 | 1 |
| 18 | 10 | 11 | 1 |
| 19 | 10 | 31 | 1 |
| 20 | 11 | 12 | 1 |
| 21 | 11 | 13 | 2 |
| 22 | 12 | 32 | 1 |
| 23 | 14 | 17 | ar |
| 24 | 14 | 21 | ar |
| 25 | 14 | 27 | 1 |
| 26 | 15 | 33 | 1 |
| 27 | 16 | 24 | 1 |
| 28 | 17 | 18 | ar |
| 29 | 17 | 34 | 1 |
| 30 | 18 | 19 | ar |
| 31 | 18 | 22 | 1 |
| 32 | 19 | 20 | ar |
| 33 | 19 | 23 | 1 |
| 34 | 20 | 21 | ar |
| 35 | 20 | 35 | 1 |
| 36 | 21 | 36 | 1 |
| 37 | 22 | 25 | 1 |
| 38 | 23 | 45 | 1 |
| 39 | 24 | 37 | 1 |
| 40 | 24 | 38 | 1 |
| 41 | 24 | 39 | 1 |
| 42 | 25 | 40 | 1 |
| 43 | 25 | 41 | 1 |
| 44 | 25 | 42 | 1 |
| 45 | 26 | 27 | 2 |
| 46 | 26 | 43 | 1 |
| 47 | 27 | 44 | 1 |
| 48 | 45 | 46 | 1 |
| 49 | 45 | 47 | 1 |
| 50 | 45 | 48 | 1 |
| 51 | 48 | 49 | 1 |
| 52 | 48 | 50 | 1 |
| 53 | 48 | 51 | 1 |

```
   54    51    52 1
   55    51    53 1
   56    51    54 1
   57    54    55 1
   58    54    56 1
   59    54    57 1
@<TRIPOS>SUBSTRUCTURE
    1 CBU        1 TEMP            0 ****  ****    0 ROOT
```

***cis*-CCBu**

```
@<TRIPOS>MOLECULE
CCB
   57    59    1    0    0
SMALL
resp

@<TRIPOS>ATOM
     1 C1B          6.3640    -1.4860    -0.9470 ca         1 CCB     -0.261883
     2 C2B          7.2400    -0.4070    -0.9230 ca         1 CCB      0.286293
     3 C3B          6.7980     0.8350    -0.4160 ca         1 CCB      0.185719
     4 C4B          5.5040     0.9800     0.0570 ca         1 CCB     -0.249740
     5 C5B          4.6150    -0.1170     0.0440 ca         1 CCB     -0.067866
     6 C6B          5.0610    -1.3420    -0.4690 ca         1 CCB     -0.169670
     7 C7B          3.2480     0.0310     0.5680 cc         1 CCB      0.452886
     8 O1B          2.4270    -0.9880     0.1690 os         1 CCB     -0.355544
     9 C8B          1.1100    -1.0280     0.5630 cc         1 CCB      0.614150
    10 C9B          0.5920    -0.0630     1.3630 cd         1 CCB     -0.671133
    11 C10B         1.3960     1.0690     1.8240 c          1 CCB      0.864095
    12 C11B         2.7830     1.0260     1.3660 cd         1 CCB     -0.660296
    13 O2B          0.9490     1.9660     2.5440 o          1 CCB     -0.633829
    14 C12B        -2.0440    -1.7910    -0.1780 ca         1 CCB      0.020970
    15 O3B          8.5090    -0.5400    -1.3890 oh         1 CCB     -0.562622
    16 O4B          7.7570     1.8170    -0.4660 os         1 CCB     -0.287542
    17 C13B        -3.2590    -2.4840     0.0360 ca         1 CCB     -0.308653
    18 C14B        -4.4850    -1.8320     0.0090 ca         1 CCB      0.266331
    19 C15B        -4.5300    -0.4400    -0.2650 ca         1 CCB      0.313400
    20 C16B        -3.3360     0.2410    -0.5070 ca         1 CCB     -0.270120
    21 C17B        -2.1090    -0.4230    -0.4620 ca         1 CCB     -0.216038
    22 O5B         -5.6940    -2.4250     0.2210 os         1 CCB     -0.336797
    23 O6B         -5.7670     0.1180    -0.2770 os         1 CCB     -0.464959
```

```
    24 C18B          7.4260     3.1080     0.0270 c3         1 CCB      -0.133058
    25 C19B         -5.7130    -3.8140     0.5020 c3         1 CCB       0.013403
    26 C20B          0.4880    -2.2580     0.0750 ce         1 CCB      -0.486439
    27 C21B         -0.8040    -2.5770    -0.1660 cf         1 CCB       0.018797
    28 H1B           6.7160    -2.4340    -1.3400 ha         1 CCB       0.200816
    29 H2B           5.1650     1.9430     0.4180 ha         1 CCB       0.171521
    30 H3B           4.3880    -2.1900    -0.4880 ha         1 CCB       0.166696
    31 H4B          -0.4400    -0.1230     1.6840 ha         1 CCB       0.251707
    32 H5B           3.4440     1.8130     1.7100 ha         1 CCB       0.225551
    33 H6B           8.9430     0.3250    -1.2820 ho         1 CCB       0.406071
    34 H7B          -3.2200    -3.5490     0.2360 ha         1 CCB       0.171459
    35 H8B          -3.3520     1.3010    -0.7340 ha         1 CCB       0.153447
    36 H9B          -1.2030     0.1340    -0.6690 ha         1 CCB       0.191766
    37 H10B          8.3240     3.7160    -0.0940 h1         1 CCB       0.109454
    38 H11B          6.6030     3.5520    -0.5470 h1         1 CCB       0.109454
    39 H12B          7.1500     3.0680     1.0880 h1         1 CCB       0.109454
    40 H13B         -6.7640    -4.0740     0.6440 h1         1 CCB       0.066103
    41 H14B         -5.3040    -4.4010    -0.3310 h1         1 CCB       0.066103
    42 H15B         -5.1540    -4.0500     1.4170 h1         1 CCB       0.066103
    43 H16B          1.2180    -3.0490    -0.0890 ha         1 CCB       0.193890
    44 H17B         -0.9630    -3.6260    -0.4140 ha         1 CCB       0.117896
    45 C22B         -5.8860     1.5170    -0.5380 c3         1 CCB       0.291978
    46 H18B         -5.3110     2.0860     0.2080 h1         1 CCB       0.003779
    47 H19B         -5.4720     1.7480    -1.5300 h1         1 CCB       0.003779
    48 C23B         -7.3640     1.8750    -0.4710 c3         1 CCB      -0.097664
    49 H20B         -7.9070     1.2610    -1.2010 hc         1 CCB       0.050846
    50 H21B         -7.7480     1.5980     0.5190 hc         1 CCB       0.050846
    51 C24B         -7.6210     3.3640    -0.7390 c3         1 CCB       0.094451
    52 H22B         -7.2230     3.6330    -1.7270 hc         1 CCB      -0.014173
    53 H23B         -7.0630     3.9690    -0.0100 hc         1 CCB      -0.014173
    54 C25B         -9.1070     3.7290    -0.6720 c3         1 CCB      -0.181897
    55 H24B         -9.2630     4.7960    -0.8660 hc         1 CCB       0.044961
    56 H25B         -9.6870     3.1660    -1.4130 hc         1 CCB       0.044961
    57 H26B         -9.5260     3.5050     0.3160 hc         1 CCB       0.044961

@<TRIPOS>BOND
     1     1     2 ar
     2     1     6 ar
     3     1    28 1
     4     2     3 ar
     5     2    15 1
     6     3     4 ar
```

| | | | |
|---|---|---|---|
| 7 | 3 | 16 | 1 |
| 8 | 4 | 5 | ar |
| 9 | 4 | 29 | 1 |
| 10 | 5 | 6 | ar |
| 11 | 5 | 7 | 1 |
| 12 | 6 | 30 | 1 |
| 13 | 7 | 8 | 1 |
| 14 | 7 | 12 | 2 |
| 15 | 8 | 9 | 1 |
| 16 | 9 | 10 | 2 |
| 17 | 9 | 26 | 1 |
| 18 | 10 | 11 | 1 |
| 19 | 10 | 31 | 1 |
| 20 | 11 | 12 | 1 |
| 21 | 11 | 13 | 2 |
| 22 | 12 | 32 | 1 |
| 23 | 14 | 17 | ar |
| 24 | 14 | 21 | ar |
| 25 | 14 | 27 | 1 |
| 26 | 15 | 33 | 1 |
| 27 | 16 | 24 | 1 |
| 28 | 17 | 18 | ar |
| 29 | 17 | 34 | 1 |
| 30 | 18 | 19 | ar |
| 31 | 18 | 22 | 1 |
| 32 | 19 | 20 | ar |
| 33 | 19 | 23 | 1 |
| 34 | 20 | 21 | ar |
| 35 | 20 | 35 | 1 |
| 36 | 21 | 36 | 1 |
| 37 | 22 | 25 | 1 |
| 38 | 23 | 45 | 1 |
| 39 | 24 | 37 | 1 |
| 40 | 24 | 38 | 1 |
| 41 | 24 | 39 | 1 |
| 42 | 25 | 40 | 1 |
| 43 | 25 | 41 | 1 |
| 44 | 25 | 42 | 1 |
| 45 | 26 | 27 | 2 |
| 46 | 26 | 43 | 1 |
| 47 | 27 | 44 | 1 |
| 48 | 45 | 46 | 1 |

```
   49    45    47 1
   50    45    48 1
   51    48    49 1
   52    48    50 1
   53    48    51 1
   54    51    52 1
   55    51    53 1
   56    51    54 1
   57    54    55 1
   58    54    56 1
   59    54    57 1
@<TRIPOS>SUBSTRUCTURE
     1 CCB         1 TEMP              0 ****  ****    0 ROOT
```

### RMSD time series for all the MD simulations

To evaluate where systems reach an equilibrium state, we report the root mean square deviation (RMSD) for the systems containing 1/2/5/10 *trans-CCBu* (Fig. 1). After equilibration, the membrane remained generally stable over the whole laps of the simulation with the RMSD oscillating around 4±1 Å.

a) 1 *trans*- molecule system:

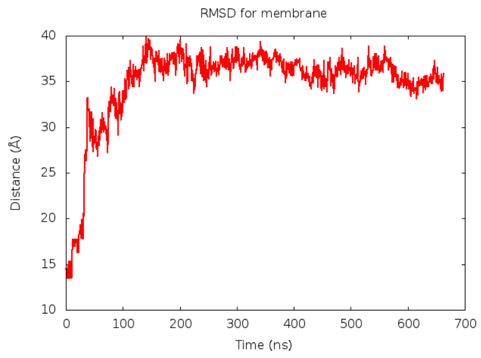

b) 2 *trans*- molecules system:

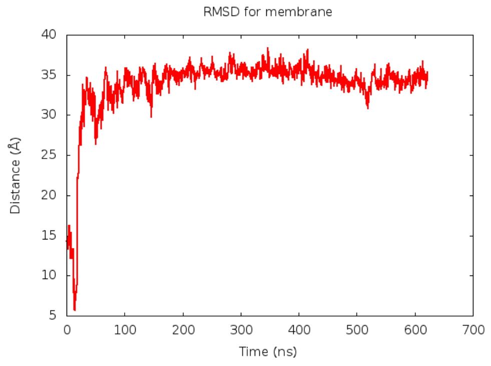

c) 5 *trans*- molecules system:

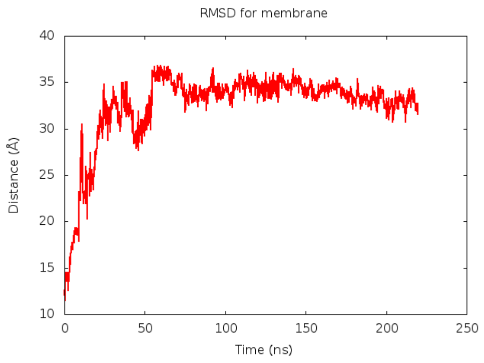

d) 10 *trans*- molecules system:

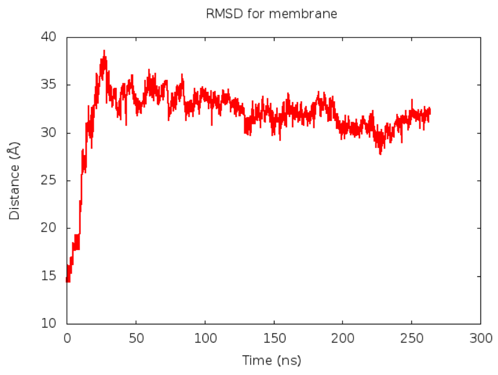

Figure S1. RMSD plots for the membrane in the system with a) 1 *trans*-; b) 2 *trans*-; c) 5 *trans*-; d) 10 *trans*-CCBu.

**Snapshots of different CCBu concentrations in both *cis-* and *trans-* conformations**

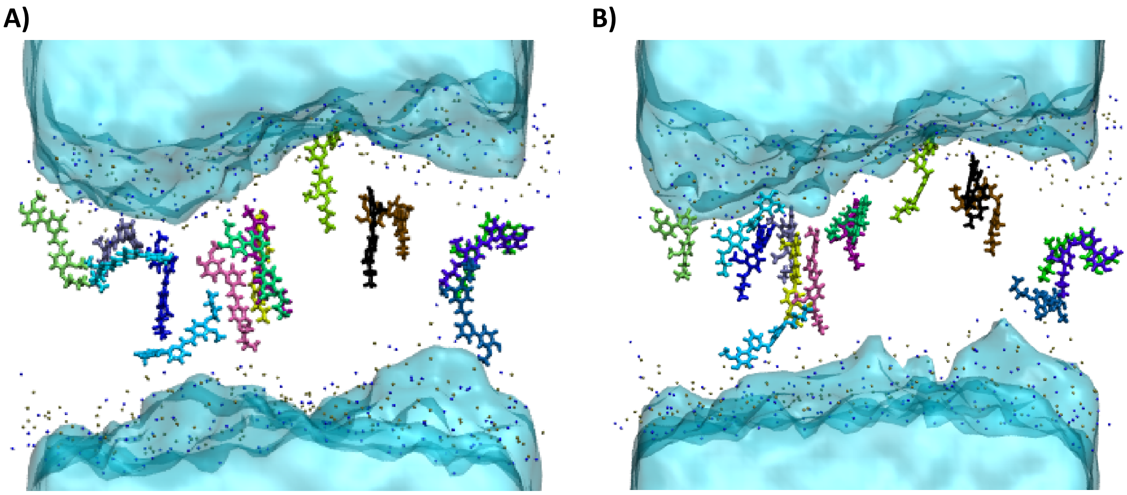

Figure S2. Representative snapshots of the MD simulations for the systems containing (A) 15 *trans* and (B) 15 *cis* CCBu chromophores embedded inside the lipid bilayer.

**Table of MD simulation time for all studied systems**

Table S1.) MD simulation time for all systems built in the study. Note that the 1 cis CCBu run was aborted once the molecule spontaneously entered the lipid bilayer, since we then switched to an enhanced sampling strategy to obtain the free energy penetration of the chromophore. Afterwards, we calculated free energy profile to be sure that the molecule will remain inside and confirm that it is its global minimum of energy.

| | **Unloaded membrane** | **1 *E-*** CCBu | **2 *E-*** CCBu | **5 *E-*** CCBu | **5 *Z-*** CCBu | **10 *E-*** CCBu |
|---|---|---|---|---|---|---|
| **MD time** | 600 ns | 660 ns | 622 ns | *660 ns | **660 ns | *660 ns |

| | **10 *Z-*** CCBu | **15 *E-*** CCBu | **15 *Z-*** CCBu | **20 *E-*** CCBu | **20 *E-*** CCBu |
|---|---|---|---|---|---|
| **MD time** | **660 ns | *660 ns | **660 ns | *660 ns | **660 ns |

***MD time after performing steered MD (SMD) to pull chromophores inside the lipid bilayer**

**** MD time after performing SMD to change the dihedral angle from *trans-* to *cis-* for chromophores**

### $-S_{CD}$ parameters for *Z* and *E* isomers

a)

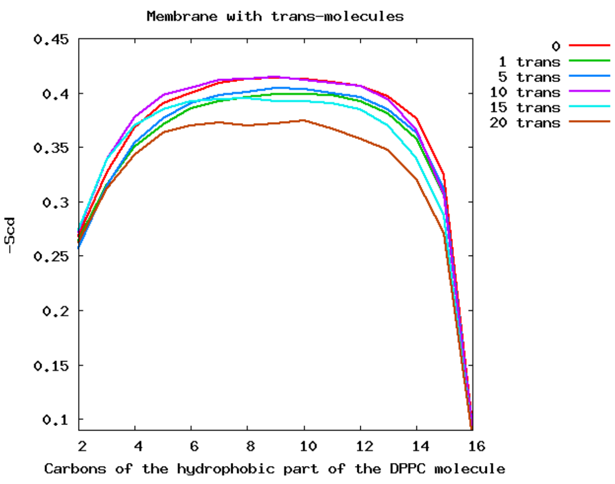

b)

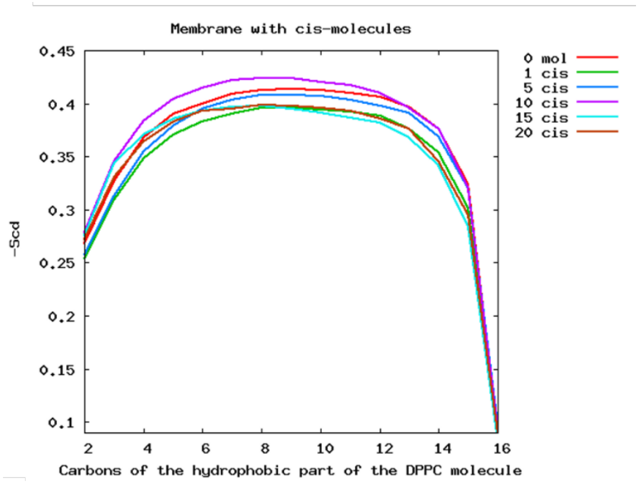

c)

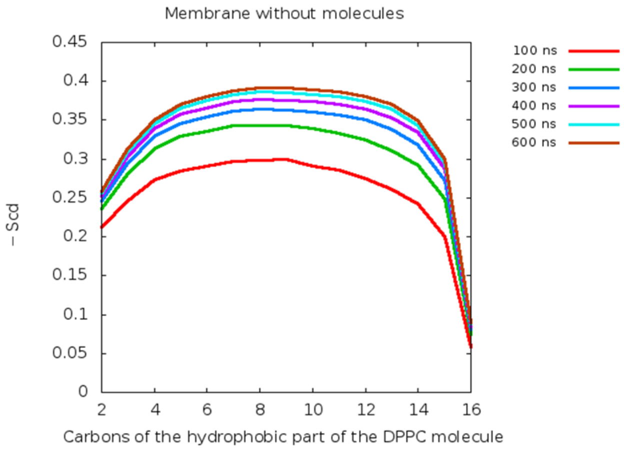

Figure S3. Average $-S_{CD}$ order parameters of DPPC tails in systems with CCBu derivatives in a) *trans*-conformation; b) *cis*-conformation, and c) time evolution of the order parameter for the lipid bilayer without any drug.

**Area per lipid for mixed E/Z CCBu loaded membranes**

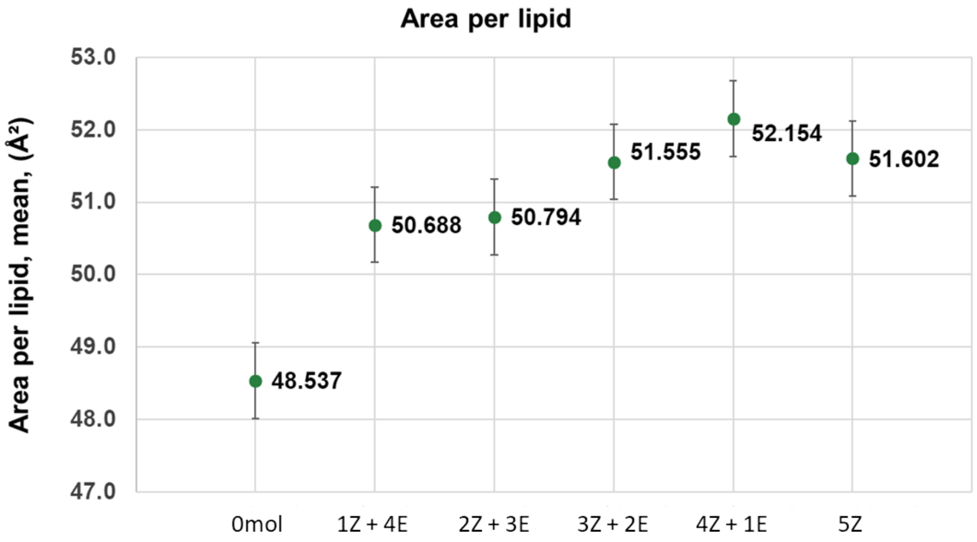

Figure S4. Average area per lipid for mixtures of E and Z isomers.

### Potential of Mean Force for the dissociation of one CCBu unit from a 5-membered aggregate

After 5-membered CCBu aggregate was firstly placed in a water box of 90x89x121Å$^3$ and equilibrated. Afterwards, the potential of mean force is estimated using meta-eABF considering the distance between the centers of mass of the leaving CCBu and the one of its original $\pi$-stacked chromophore remaining in the aggregate. The collective variable is explored in the range between 3.0 and 20 Å. Note that to simplify the conformational landscape to be the distance between the 4-remaining CCBu units have been constrained to 3 Å

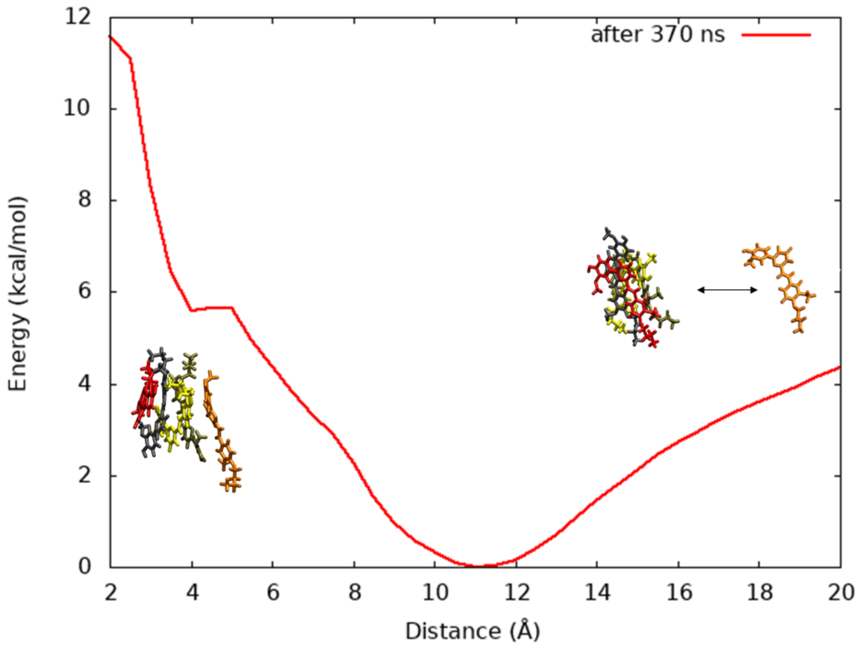

Figure S5. Potential of Mean force for the dissociation of one CCBu form a 5-membered aggregate.
